# Identification of natural products and synthetic analogs which inhibit microsporidia spores and prevent infection

**DOI:** 10.1101/2025.08.06.669004

**Authors:** Qingyuan Huang, Lauren E. Brown, Guoqing Pan, Junhong Wei, John A. Porco, Jie Chen, Aaron W. Reinke

## Abstract

Microsporidiosis, a disease caused by microsporidia, affects many animals, with symptoms ranging from diarrhea to death, especially in immunocompromised individuals. Current treatments, such as albendazole and fumagillin, are limited in efficacy. To address this problem, we utilized *Caenorhabditis elegans* infected with its natural microsporidian *Nematocida parisii* to evaluate 4,080 structurally diverse compounds from the Boston University Center for Molecular Discovery (BU-CMD) chemical library. From this screen we identified 34 compounds that effectively inhibited *N. parisii* infection and restored the reproductive capacity of *C. elegans*. All 17 compounds we validated prevented *N. parisii* infection in *C. elegans*, and 10 were capable of suppressing microsporidia invasion by inactivating mature spores. Additionally, five of the identified compounds were also effective against *Pancytospora epiphaga*, a species related to human*-*infecting microsporidia. Together this work identifies and characterizes compounds which inhibit microsporidia infection.

**Importance:** Microsporidia are a large group of microbial parasites that infect many animals, including humans. Many agriculturally important animals such as honey bees, shrimp, crabs, and fish are infected by microsporidia, and these infections often result in smaller animals, fewer offspring, and death. Only two drugs are commonly used to treat microsporidia infections; due to some species being resistant and host toxicity there is a need to identify other microsporidia inhibitors. In this study, we screened over 4000 small molecules using a model round worm infected with microsporidia. We identified several dozen inhibitors and characterized how these compounds prevented infection by determining which stage of the parasite they act upon. Together our work identified and characterized compounds that could be used as a starting point to design better microsporidia inhibitors.

## Introduction

Microsporidiosis, a zoonotic disease, has led to significant health issues among high-risk populations, particularly those with compromised immune systems (Taghipour et al. 2022). When mature spores are shed through the feces of infected hosts, they can remain infective for a long time in the environment (Dunn and Smith 2001). The majority of parasite infections in animals occur via the fecal-oral route through contaminated food or water (Javanmard et al. 2018; Taghipour et al. 2022). Immunocompetent individuals are usually asymptomatic, whereas immunocompromised individuals suffer from prolonged diarrhea as a major consequence of microsporidia infection (Han et al. 2021). Alarmingly, it is difficult to treat these infections due to the lack of knowledge of microsporidia-specific cellular targets, resistant microsporidia species, and a lack of effective anti-microsporidia drugs. In addition to humans, microsporidia infect many types of animals, including those with economic and food security importance such as honey bees, shrimp, and silkworms (Stentiford et al. 2016; Murareanu et al. 2021). There is a critical need to develop effective and safe drugs for treating microsporidia infections, especially in chronically ill individuals and economically valuable animals.

There are currently no effective drugs that can treat all species of microsporidia without causing adverse side effects. Benzimidazole analogs, which inhibit microsporidia proliferation, are one of the most effective classes of compounds to control microsporidiosis, for example, using albendazole against shrimp infected with *Enterocytozoon hepatopenaei* (Subash et al. 2023). FangWeiLing (FWL), a carbendazim-based compound, is the only pharmaceutical approved in China for pébrine disease, which is caused by microsporidia infection of silkworms (Xing et al. 2022). However, silkworms are also adversely affected by high concentrations of benzimidazoles (Jyothi and Patil). In addition, albendazole only has a limited effect on microsporidiosis caused by the human-infecting species *Encephalitozoon bieneusi* and *Vittaforma cornea*, which encode beta-tubulin associated with albendazole resistance (Karin Leder, Norbert Ryan, Denis Sp 1998; Akiyoshi et al. 2007; Franzen and Salzberger 2008). In contrast with albendazole, the antimicrobial natural product fumagillin is effective in the treatment of *E. bieneusi* (Molina et al. 2000). However, fumagillin has been shown to have toxic side effects such as causing reversible thrombocytopenia as well as aseptic meningitis in humans (Wei et al. 2022). Other types of inhibitors that act on spores to prevent infection have been identified such as flavone analogs which inhibit microsporidia by preventing spore germination and benzenesulfonamide derivatives which inhibit microsporidia infection by inactivating spores (Huang, Chen, Pan, et al. 2023; Huang et al. 2025). However, the safety and efficacy of these compounds in treatment of microsporidiosis has not yet been confirmed. Therefore, it is necessary to identify additional microsporidia inhibitors.

The Boston University Center for Molecular Discovery (BU-CMD) screening collection is a curated library of academically-derived compounds created through both diversity-oriented and target-oriented synthesis methods, and is also replete with a large number of synthetic analogs and intermediates of bioactive natural products (Brown et al. 2011). The collection aims to achieve a higher level of structural complexity and a wider range of chemical space than traditional combinatorial chemistry libraries (Manier et al. 2017). This library has been used to discover inhibitors of many kinds of pathogenic microbes including the yeast *Candida auris*, the protist *Leishmania*, and orthopoxviruses (Dower et al. 2012; Iyer et al. 2020; Kavouris et al. 2023).

The model organism *Caenorhabditis elegans* and its natural microsporidian parasite *Nematocida parisii* are a convenient system in which to identify microsporidia inhibitors (Murareanu et al. 2022; Huang, Chen, Pan, et al. 2023; Huang et al. 2025). Infection of *C. elegans* begins when the worms ingest spores, which then germinate in the intestinal lumen, depositing a sporoplasm inside an intestinal cell. The sporoplasm then proliferates in the intestinal cells as meronts, which differentiate into spores, that then exit the animal (Gang and Lažetić 2024). Infection of *C. elegans* causes smaller animals with reduced progeny, and this phenotype can be used in liquid assays in 96-well plates to identify compounds which reverse the detrimental effects of infection (Balla et al. 2016).

To identify novel microsporidia inhibitors, we screened the BU-CMD’s chemical library, containing 4,080 compounds at the time of screening. By utilizing the highly effective *C. elegans*-*N. parisii* system, we identified 34 compounds that restored the ability of *C. elegans* to produce progeny in the presence of *N. parisii*. We validated 17 of the compounds, showing that all of them could reduce *N. parisii* infection levels. Several of the compounds inactivated *N. parisii* spores and prevented invasion of *C. elegans*. We also tested five of the validated inhibitors and all of them inhibited *Pancytospora epiphaga*, a microsporidia species related to those that infect humans. All together we identified many microsporidia inhibitors, providing a starting point for the design of more potent inhibitors based on these scaffolds.

## Results

### BU-CMD chemical library screening identifies microsporidia inhibitors

To identify compounds from the BU-CMD chemical library that inhibit microsporidia infection, we utilized a previously described 96-well infection assay (Murareanu et al. 2022; Huang, Chen, Pan, et al. 2023; Huang et al. 2025). *N. parisii* spores were incubated with BU-CMD compounds, and after one hour, *C. elegans* at the L1 larval stage were added. The cultures were then maintained on a shaker at 21°C for 5 days. Since *N. parisii* infection reduces the reproductive capacity of *C. elegans*, the effectiveness of the inhibitors was assessed by counting worm offspring (Balla et al. 2016; Willis, Zhao, Sukhdeo, Wadi, Jarkass, et al. 2021). Worms were then stained with rose bengal, imaged using a flatbed scanner, and counted using automated image analysis (Murareanu et al. 2022). We screened 4,080 compounds from the BU-CMD library, each at a concentration of 60 µM. Each compound was screened in triplicate, resulting in the quantification of over 12,000 interactions. Our screen was judged to be reliable at distinguishing true interactions with a Z-factor of 0.46 (Data S1). Compared to uninfected controls, 34 compounds increased progeny production in infected worms by 60% (Fig. 1A and Fig. S1A). Compared to infected controls, these 34 compounds increased progeny production 2.6-6.3-fold (Fig. S1B and Data S1).

**Fig 1.**
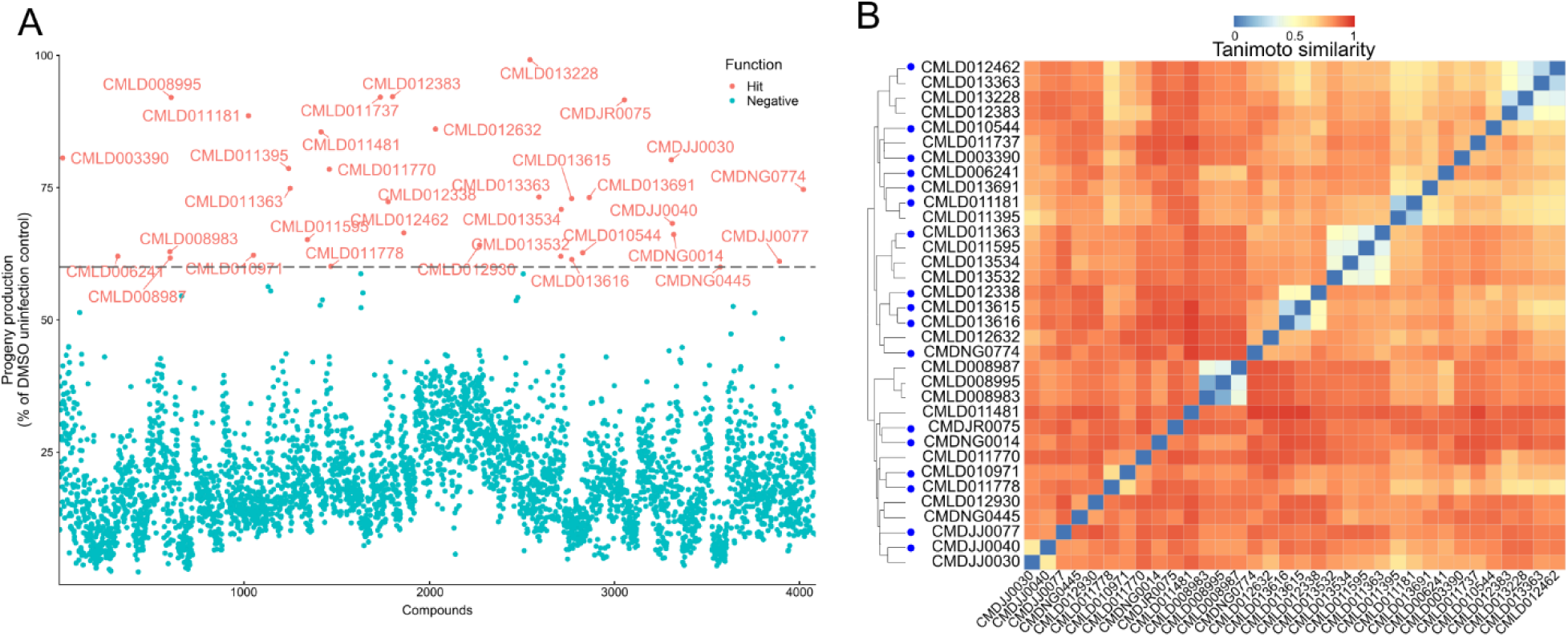
High-throughput screen of 4,080 BU-CMD compounds identifies inhibitors of *N. parisii*. **(A)** *N. parisii* spores were incubated with compounds for 1 hour, after which L1 stage *C. elegans* were added and incubated for five days. Progenies were then stained and quantified. Each point represents the mean progeny number as a percentage of the uninfected control. Compounds with at least 60% progeny production are colored red and their compound identification labels are shown. Compounds with less than 60% progeny production are colored blue. Three independent biological replicates were performed for each compound. **(B)** Compounds were clustered by compound similarity and displayed as a heat map according to the scale with 0 (blue) being the most similar and 1 (red) being the least similar. Compounds tested in subsequent experiments are indicated by dark blue dots next to the compound name.

The structural similarity of the candidate inhibitors was analyzed using hierarchical clustering (Fig. 1B and Fig. S2). We identified several clusters of synthetic heterocycles, including pyridopyrimidinones (CMLD011363, CMLD011595, CMLD013534, and CMLD013532) previously shown to espouse anti-poxviral activity, and bis-phenolic diarylated indoles (CMLD012338, CMLD013615, and CMLD013616) resembling the phenisatin class of laxatives (Dadashpour and Emami 2018; Brown et al. 2022). Several identified compounds were either natural products or synthetic analogs of established bioactive natural product classes. For example, we identified the xanthone natural product garcinexanthone C (CMLD013228) (Chen et al. 2008) alongside several closely-related synthetic xanthonoid analogues (CMLD012462, CMLD013363 and CMLD012383), as well as one iodinated tetrahydroxanthone derivative (CMLD010971) resembling the natural product blennolide B (Zhang et al. 2008; Qin et al. 2015). Two chalconoid-type compounds, close congeners of the natural product nicolaioidesin C, were also identified (CMLD011181 and CMLD011395) (Cong et al. 2008a; Laflamme et al. 2025). Interestingly, three of the hits (CMLD008995, CMLD008987, and CMLD008983) were synthetic compounds obtained *via* structural “remodelling” of fumagillol, the hydrolysis product of fumagillin (Balthaser et al. 2011). However, upon confirmatory quality control analysis by NMR and LC/MS, we found that the stocks of these hits were contaminated with varying amounts of fumagillol, which likely explains the microsporidia inhibition observed. Other singleton hits were identified as closely related to members of the azaphilone (CMLD003390) (Achard et al. 2012), and epoxyquinol (CMLD011481) (Bardhan et al. 2006) natural product families and a synthetic precursor to the natural product sanggenol F (CMLD010544) (Qi et al. 2016) which was also identified. Lastly, we also identified two additional natural product singletons, curcumin (CMLD011737, also previously identified in a 2022 screen) and glabridin (CMLD013691), as inhibitors (Murareanu et al. 2022). We ultimately selected 17 compounds which represent a diversity of chemical structures to validate, prioritizing those which were less likely to be pan-assay interference (PAINS) compounds (Baell and Holloway 2010).

### Validation of BU-CMD compounds which inhibit *N. parisii* infection

To assess whether the inhibitor candidates directly affect microsporidia infection, we conducted continuous infection assays, following a protocol similar to the initial screening. We incubated *N. parisii* spores with the inhibitors in 24-well plates for 1 hour, followed by the addition of L1 stage worms, which were cultured for 4 days. We then fixed the worms and stained with Direct Yellow 96 (DY96) which binds to the chitin in the microsporidia spores and the embryos from *C. elegans* (Willis et al. 2022). We included the known microsporidia inhibitor dexrazoxane as a positive control (Murareanu et al. 2022). All inhibitors increased the proportion of gravid nematodes (containing at least one embryo) within infected *C. elegans* populations (Fig. 2A). These results are consistent with our screen showing that these compounds can rescue the ability of *C. elegans* to produce progeny in the presence of *N. parisii*. A comparison of embryo counts from our validation experiments to quantified progeny production from the screen shows a weak correlation (Fig S3). To determine if the compounds could directly prevent infection, we quantified the proportion of the population that generated *N. parisii* spores. We observed that 15 of the inhibitors significantly reduced the formation of newly matured spores (Fig. 2B). The other two inhibitors, CMDNG0774 and CMLD011778, showed relatively weak microsporidia inhibition. To determine if these two compounds had any impact on *N. parisii* infection, we categorized infection based on severity and found that both compounds significantly reduced the levels of severe infection (Fig. 2C-E). Overall, all 17 compounds we validated demonstrated the ability to inhibit microsporidia infection.

**Figure 2.**
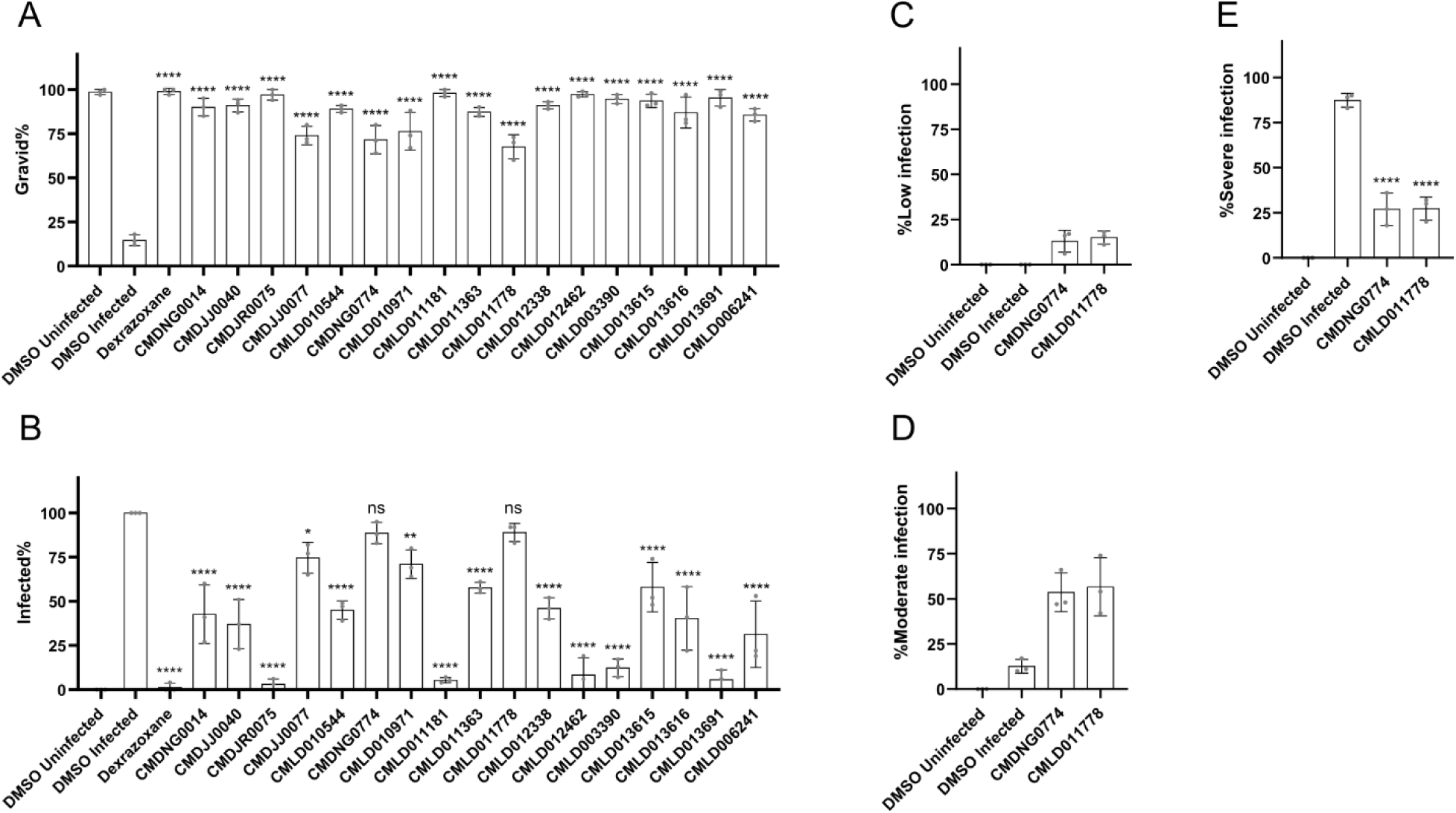
Validation that BU-CMD compounds inhibit *N. parisii*. **(A-E)** *N. parisii* spores were incubated with compounds for one hour and then L1 stage N2 worms were added and cultured for 4 days. Worms were then fixed and stained with DY96, which labels *N. parisii* spores and *C. elegans* embryos. **(A)** Quantification of the percentage of gravid animals (those which contain embryos). **(B)** Quantification of the percentage of infected worms (those containing newly formed spores). **(C)** Quantification of worms displaying low infection (defined as less than half of an animal with spores). **(D)** Quantification of worms displaying moderate infection (defined as half an animal with spores). **(E)** Quantification of worms displaying high infection (defined as both halves of an animal with spores). n= 3 biological replicates, N = ≥ 100 worms counted per biological replicate. The P-values were determined by one-way ANOVA with post hoc test, with all comparisons to DMSO-infected controls. Means ± SD (horizontal bars) are shown. (*p <0.05, **p < 0.01, ****p < 0.0001, ns means not significant).

### None of the identified compounds inhibit microsporidia proliferation

A key strategy for preventing microsporidian infection is to inhibit the intracellular form of the parasite, blocking proliferation and subsequent spore formation. To test if the validated compounds could inhibit proliferation, we infected worms with *N. parisii*, washed away undigested spores, and incubated the worms with the BU-CMD compounds or dexrazoxane in 24-well plates. Only the control compound dexrazoxane significantly increased the proportion of gravid worms (Fig. 3A). None of the BU-CMD compounds displayed a decrease in the proportion of the population generating spores, under conditions where inhibition by dexrazoxane was observed (Fig. 3B). To determine if there is a reduction in parasite at an earlier stage of development, we performed the experiment as described above, except we fixed worms at either 2 days (before new spore formation) and 4 days (after new spore formation) post infection, and stained meronts using fluorescence in situ hybridization (FISH) with an *N. parisii* 18S RNA-specific probe (Troemel et al. 2008). At either 2 days or 4 days post infection, no compounds besides dexrazoxane showed a significant reduction in meronts (Fig. 3C and D).

**Figure 3.**
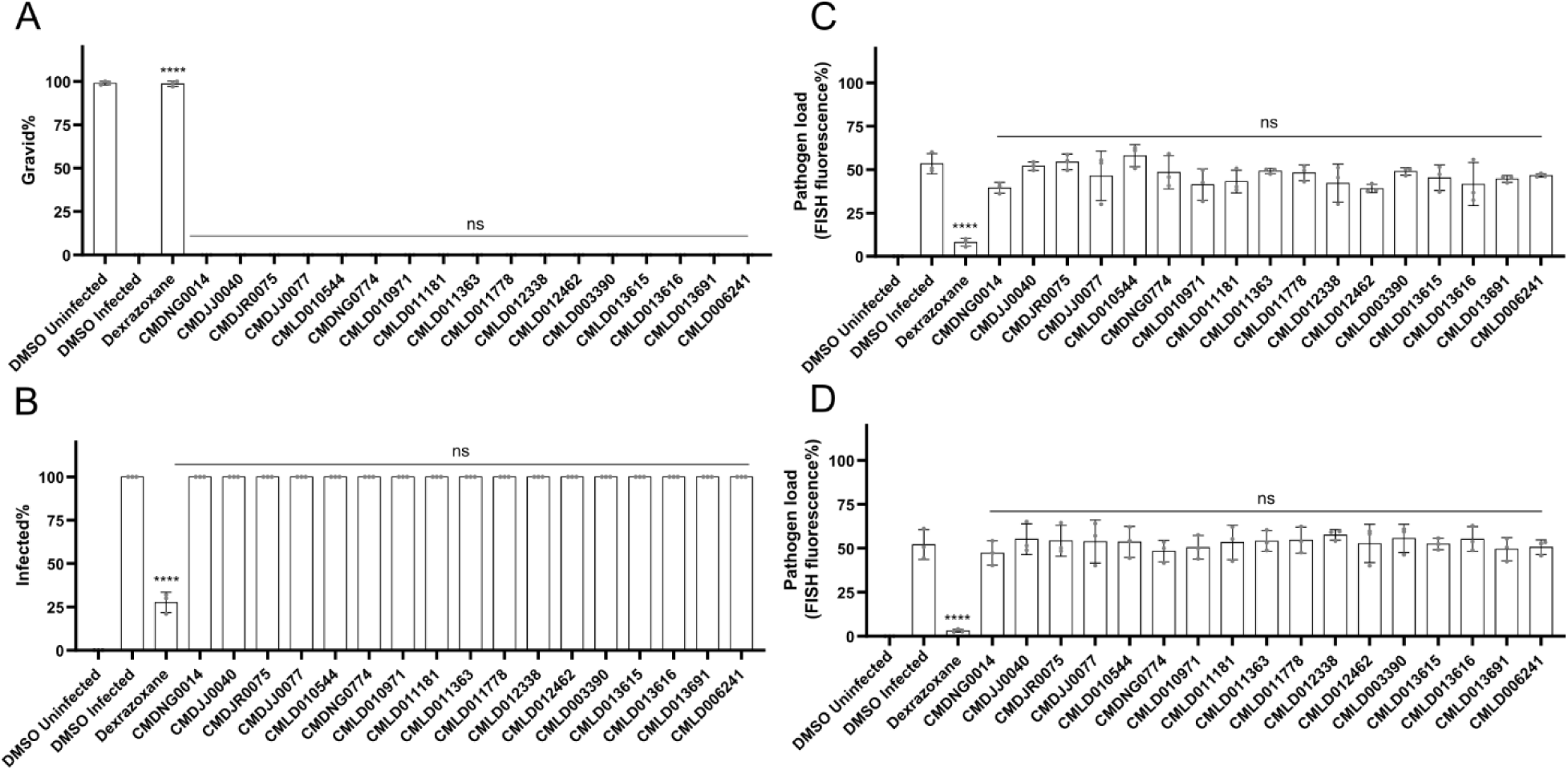
BU-CMD compounds do not inhibit microsporidia proliferation. **(A-E)** L1 stage *C. elegans* were exposed to *N. parisii* spores for 3 hours and then washed to remove undigested spores. Infected *C. elegans* were then incubated with compounds for 2 (C) or 4 (A, B, D, and E) days, fixed, and stained with DY96 and a FISH probe specific to the *N. parisii* 18S rRNA. (**A)** The percentage of gravid worms. **B** The percentage of worms that contain newly formed spores. (**C and D)** Quantification of FISH fluorescence at 2 (C) or 4 (D) days following infection. n = 3 biological replicates, N = ≥ 100 worms (A, and B) or N=10 (C, D) counted per biological replicate. The P-values were determined by one-way ANOVA with post hoc test, with all comparisons to DMSO infected, except for those indicated by brackets. Means ± SD (horizontal bars) are shown. (*p <0.05, ***p < 0.001, ****p < 0.0001, ns means not significant).

### Several BU-CMD inhibitors inactivate *N. parisii* spores and prevent invasion

The spore is the environmentally resistant extracellular form of microsporidia and inactivating the spore would prevent host invasion (Huang, Chen, Lv, et al. 2023). To determine if any of the validated BU-CMD compounds caused *N. parisii* mortality, we exposed spores to the compounds for 24 hours and used Sytox Green to specifically stain dead spores. Spore viability was assessed by counting the proportion of Sytox Green-positive spores, with acetone treatment used as a positive control for spore killing. We identified 10 inhibitors capable of significantly increasing spore mortality (Fig. 4A). After treating spores with inhibitors, we exposed them to worms and determined the ability of *N. parisii* to invade *C. elegans* by counting the number of sporoplasms. All the tested BU-CMD inhibitors caused a reduction in the sporoplasms numbers (Fig. 4B). The compounds which had amongst the greatest reduction in sporoplasm numbers (CMDJR0075, CMLD013691, CMLD012462, and CMLD011181), also caused the highest mortality to spores (Fig. 4A and B). We also assessed the ability of compounds to block spore germination. Only the control compound ZPCK, a known inhibitor of germination, prevented microsporidia germination (Fig. S4A) (Murareanu et al. 2022). Additionally, none of the compounds could trigger spore firing *in vitro* (Fig. S4B).

**Figure 4.**
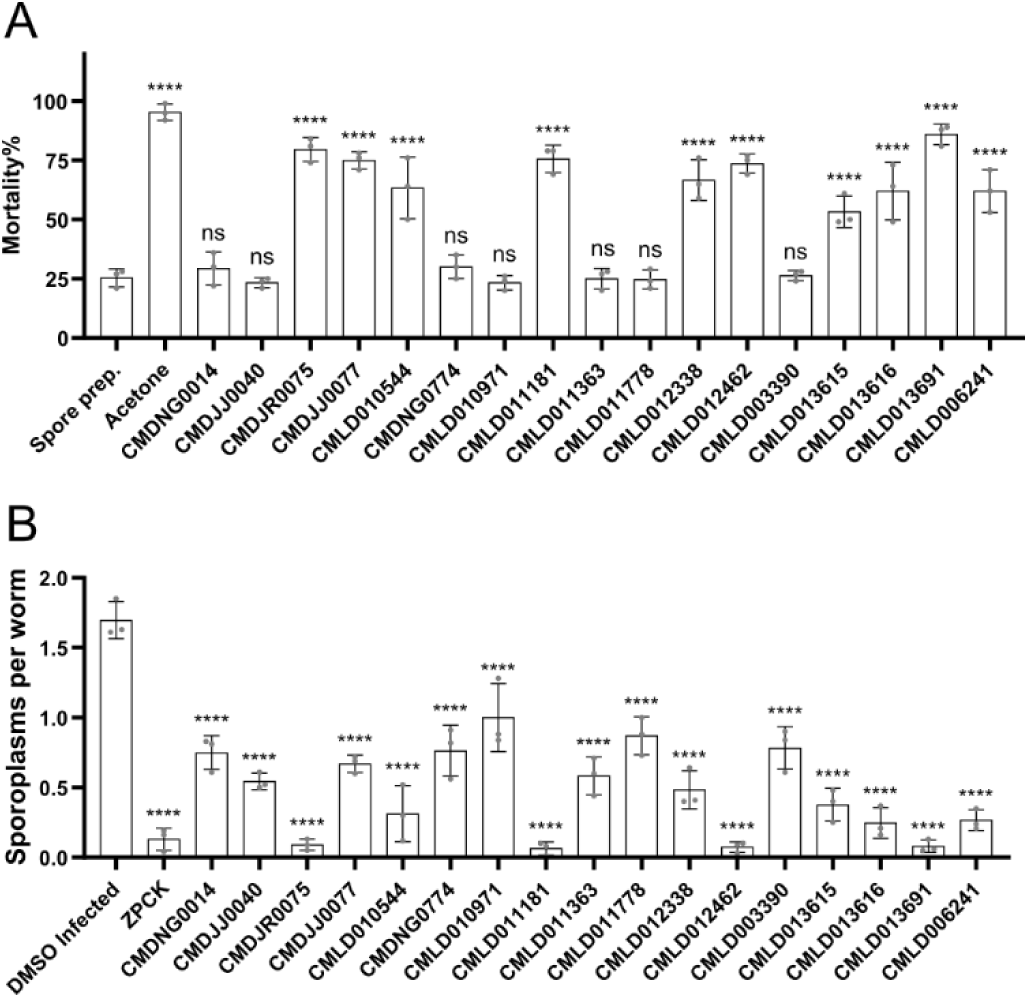
Several BU-CMD compounds inactivate *N. parisii* spores and prevent invasion of *C. elegans*. **(A)** *N. parisii* spores were treated with the indicated compounds for 24 hours, followed by Sytox Green and Calcofluor White M2R staining. Percentage of non-viable spores. **B** After incubating *N. parisii* spores with compounds for 24 hours, spores were washed and added to L1 stage worms. After incubation for 3 hours, worms were fixed, the spores (DY96) and sporoplasms (FISH) were stained. (**B)** The mean number of sporoplasms per worm. n = 3, N = ≥ 100 spores (A) or N = ≥ 100 worms (B) counted per biological replicate. The P-values were determined by one-way ANOVA with post hoc test. Means ± SD (horizontal bars) are shown. (**p < 0.01, ***p < 0.001, ****p < 0.0001, ns means not significant).

### Identified BU-CMD inhibitors inhibit *P. epiphaga*

To determine whether the *N. parisii* BU-CMD inhibitors we identified could also block infections by other microsporidia species, we chose five structurally distinct inhibitors and evaluated their effectiveness against *Pancytospora epiphaga*. This microsporidian infects the hypodermis and muscles of *C. elegans* and is part of the *Enterocytozoonida* clade, which includes notable human pathogens such as *V. cornea* and *E. bieneusi* (Zhang et al. 2016; Bojko et al. 2022). We infected *C. elegans* with *P. epiphaga* and measured the pathogen burden in the presence of dexrazoxane or each of five BU-CMD inhibitors. We observed that all five inhibitors significantly reduced *P. epiphaga* infection levels (Fig. 5).

**Figure 5.**
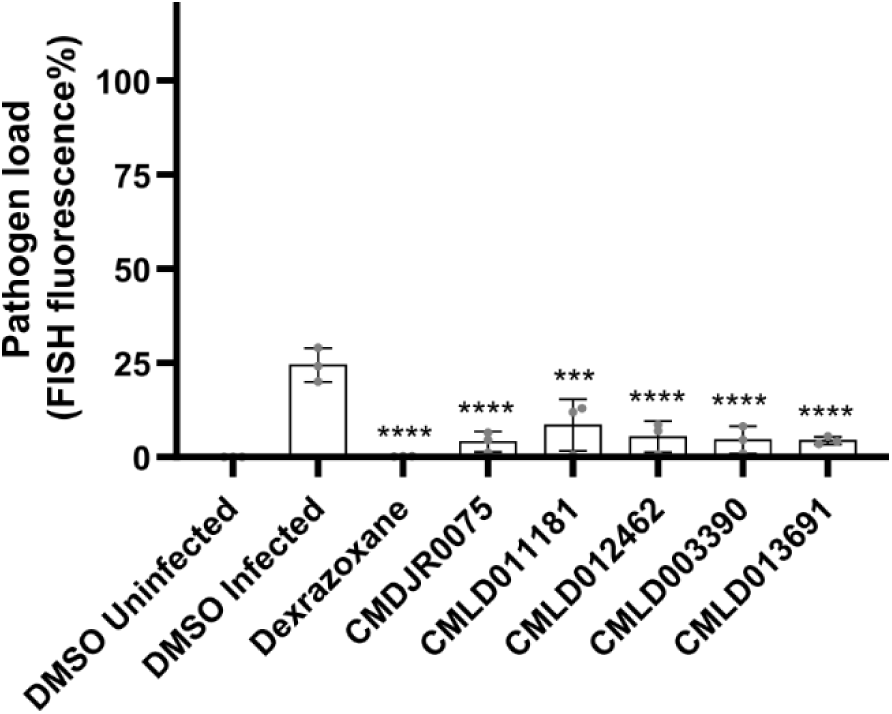
Several BU-CMD compounds inhibit *P. epiphaga* infection of *C. elegans*. **(A)** *P. epiphaga* spores were incubated with the indicated compounds for 1 hour. L1 stage *C. elegans* were then added and cultured for 4 days. Animals were then fixed and stained with a *P. epiphaga* 18S rRNA FISH probe. Quantification of pathogen load. n = 3 biological replicates, N = 10 animals. The P-values were determined by one-way ANOVA with post hoc test. Means ± SD (horizontal bars) are shown (**p < 0.001 and ****p < 0.0001).

## Discussion

In this study, we performed the largest reported screen for microsporidia inhibitors to date, assaying 4,080 compounds from the BU-CMD’s curated chemical repository for the ability to inhibit *N. parisii* infection of *C. elegans*. After PAINS filtering, we confirmed inhibitor activity for all 17 compounds carried forward for validation, representing diverse structural classes. These inhibitors prevent infection through a variety of mechanisms including causing spore mortality and blocking proliferation. We did not attempt to follow up on all the compounds from this screen with activity against *N. parisii*, and thus there are other potential inhibitors from the screen that remain to be validated and characterized.

Including the results of this study, *C. elegans* infected with *N. parisii* has now been used to screen four compound collections for a total of ∼10,000 molecules (Murareanu et al. 2022; Huang, Chen, Pan, et al. 2023; Huang et al. 2025). The hit rates from the three previously reported screens vary from 0.4-1%, which is similar to the 0.8% rate which we observe with this collection. These screens have identified multiple ways in which microsporidia can be inhibited by small molecules (Fig. 6). One mechanism is compounds acting directly on the spores, either making them inviable or preventing germination. Alternatively, the compounds can act on the intracellular stage of the parasite and prevent proliferation. Including this study, of 53 validated inhibitors, only 3 have been reported to block proliferation: dexrazoxane, the albendazole analog MMV1782387, and the benzenesulfonamide 5357859. The commonly used microsporidia inhibitor fumagillin has also been shown to block proliferation in *C. elegans* (Murareanu et al. 2022). Together the results of these screens suggest that inhibitors of proliferation are rare, which is potentially due to challenges in achieving a compound concentration high enough to penetrate both animal and microsporidian membranes without causing host toxicity. Including this study, 15 compounds found to inhibit *N. parisii* also have activity against *P. epiphaga*, demonstrating the ability of *N. parisii* to be used to find broad-spectrum microsporidia inhibitors.

**Figure 6.**
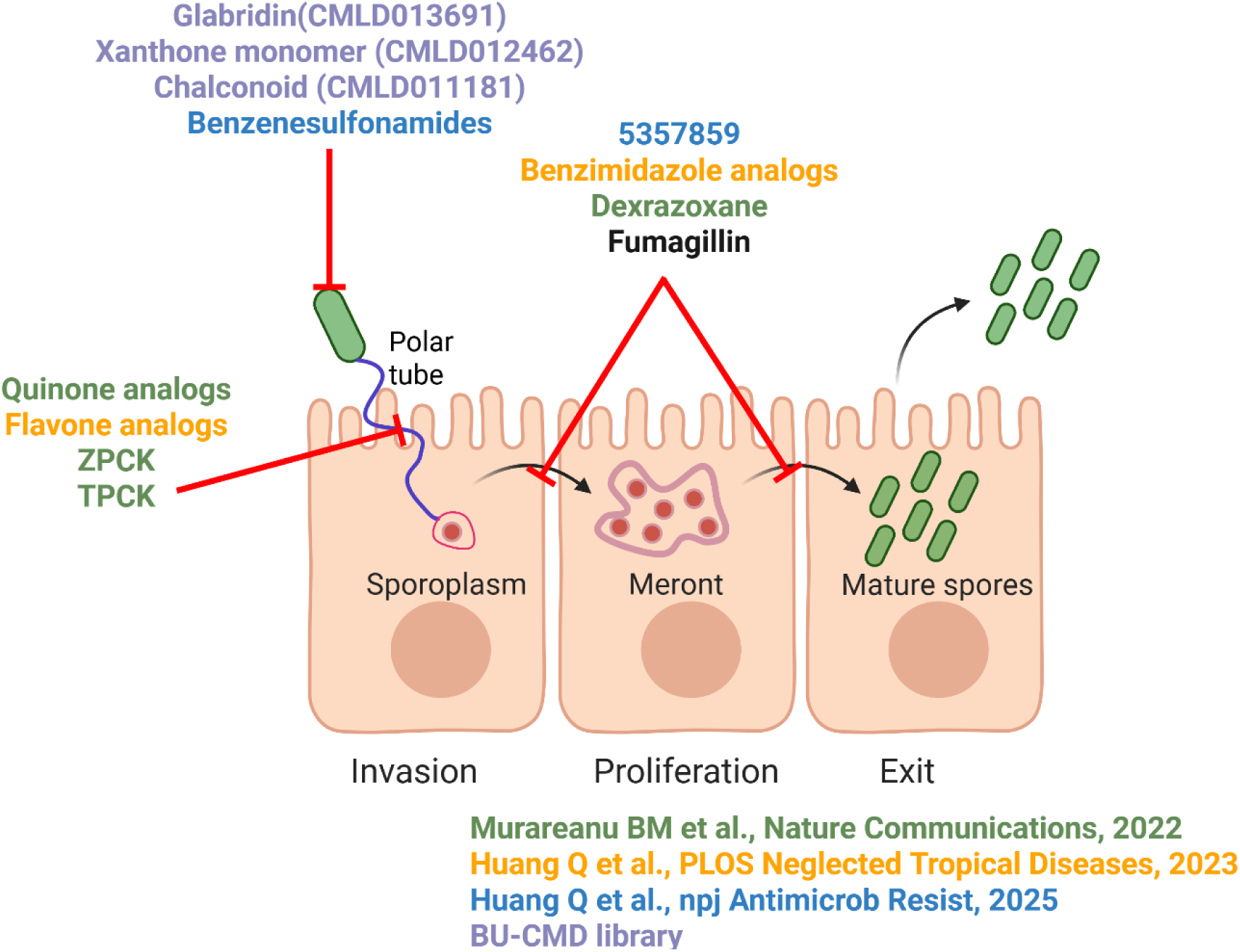
Summary of inhibitor screens using *N. parisii* and *C. elegans*. Schematic of where in the microsporidia life cycle that compounds were determined to function, either inactivating microsporidia spores, blocking the germination of spores, or preventing microsporidia proliferation. Compounds are colored based on the study that they were identified in, according to the legend at the bottom right.

The four strongest inhibitors identified in this study all function by causing spore mortality. We previously identified several microsporidia inhibitors including benzenesulfonamides which can kill spores (Huang et al. 2025). Other types of compounds that cause mortality or reduce spore numbers have been reported such as porphyrins and massetolides (Buczek et al. 2020; Jarkass et al. 2024). The strongest inhibitor we found is the benzenesulfamate ester CMDJR0075, which bears a *tert*-butylphenyl moiety (Blackburn et al. 2017). We also identified the weak inhibitor CMDNG0774, which is similar in structure to the halogenated benzenesulfonamides we previously characterized (Kelleghan et al. 2021; Huang et al. 2025). The other three inhibitors were the natural product glabridin (CMLD013691), a prenylated xanthone (CMLD012462), and a cyclohexenyl dihydrochalcone (CMLD011181). Glabridin is a flavonoid found in licorice root that has a variety of biological effects including antimicrobial activity against pathogenic bacteria and fungi (Zhang et al. 2023). Xanthones are plant-derived compounds that have been reported to elicit antimicrobial activity (Gopalakrishnan et al. 1997; Liu et al. 2022). Cyclohexenyl dihydrochalcones are a natural product derived from the Diels-Alder cycloaddition of prenylflavonoid dienes and chalcones (Cong et al. 2008b). Both chalcones and dihydrochalcones are broadly known to espouse antioxidant, anti-infective and anticancer properties (Rozmer and Perjési 2016; Nematollahi et al. 2023). These compounds that inactivate microsporidia spores could be developed into antiseptics to prevent infection by microsporidia that commonly infect important agricultural animals such as honey bees, silk worms, and shrimp (van den Heever et al. 2016; Chaijarasphong et al. 2021; Huang et al.).

## Acknowledgements

**Funding:** We thank Edward James, Yin Chen Wan, and Jonathan Tersigni for helpful comments on this manuscript. We thank the Boston University Center for Molecular Discovery for providing the BU-CMD compound library and the individual compounds to retest. This work was supported by Canadian Institutes of Health Research grant (no. 461807 to A. W. R.) and Q. H. was supported by an award from the China Scholarship Council. Work at the BU-CMD, including curation and distribution of the repository screening collection, is supported by NIH U01 TR002625. Compound contributions from Jeffrey Johnson (CMDJJ0030, CMDJJ0040, CMDJJ0077), Jennifer Roizen (CMDJR0075) and Neil Garg (CMDNG0014, CMDNG0445, CMDNG0774) and members of their respective research laboratories are gratefully acknowledged. **Competing interests:** The authors declare that they have no competing interests.

## Material and methods

### *C. elegans* maintenance

The wild-type *C. elegans* strain N2 was cultured on nematode growth media (NGM) plates and maintained using the *Escherichia coli* OP-50 strain (Lewis and Fleming 1995). To produce age-synchronized worms, M9 solution was used for washing the worms from the NGM plates, and sodium hypochlorite and sodium hydroxide were used for bleaching. The embryos were released into the solution and incubated in M9 buffer at 21 °C for 18 to 24 hours until they hatched.

### Bacterial growth

*E. coli* OP-50 from a frozen stock was streaked onto LB agar plates and incubated overnight at 37 °C. To create OP50 stocks, a single colony was inoculated into LB and incubated for 18 hours at 37 °C with shaking. Cultures were then concentrated using centrifugation to achieve a 10x concentration and then stored at 4°C.

### Microsporidia spore preparation

Stocks of *N. parisii* spores were generated using a liquid protocol that was modified from a previous plate based approach (Troemel et al. 2008; Willis, Zhao, Sukhdeo, Wadi, Jarkass, et al. 2021). ∼6000 L1 stage *C. elegans* N2 worms were infected with *N. parisii* spores (10,000 spores/μL) in 3 mL of S-complete medium containing OP-50 (Hibshman et al. 2021). The worms were incubated for 4 days at 21°C with rotation, then transferred to 50 mL of S-complete medium with OP-50 and 50,000 L1 worms which was incubated for 6 days with shaking. The infected worms were harvested and stored at -80 °C. The worms were then mechanically disrupted using 2 mm diameter zirconia beads and debris was removed by passing the homogenate through a 5 μm filter (Millipore). *N. parisii* spore preparations were verified to be free of bacterial and fungal contamination and subsequently stored at -80 °C. The concentration of spores was measured by counting DY96-stained spores using the Cell-VU sperm counting slide (Willis, Zhao, Sukhdeo, Wadi, El Jarkass, et al. 2021).

### Phenotypic screening of the BU-CMD library to identify microsporidia inhibitor

A total of 4080 compounds were provided by the Boston University Center for Molecular Discovery (BU-CMD). We screened these compounds using a previously described method to quantify the ability of compounds to improve the reproductive capacity of *C. elegans* in the presence of *N. parisii* (Murareanu et al. 2022; Huang, Chen, Pan, et al. 2023; Huang et al. 2025). We generated 96-well plates containing 100 bleach-synchronized L1 worms, 1% DMSO, 18,000 spores/μL *N. parisii,* and 60 μM test compounds. Column 1 was used for the uninfected controls which did not contain any *N. parisii* spores and column 2 was used for the infected controls which contained spores, but no inhibitor. Each well contained 5x OP50 in 50 μL of K-medium (51 mM NaCl, 32 mM KCl, 3 mM CaCl₂, 3 mM MgSO₄, 3.25 μM cholesterol). Compounds were added using a 96-well pinning tool from V&P Scientific and spores were incubated with compounds for 1 hour at ∼21 °C before adding worms. The 96-well plates were then covered with a breathable adhesive porous film, placed inside a humidity box which was wrapped in parafilm, and incubated at 21 °C with shaking at 160 rpm for 5 days. Three biological replicates were performed for each compound.

### Quantification of progeny production

After a five-day incubation period, 10 μL of a 0.3125 mg/mL Rose Bengal solution was dispensed into each well of the screening plate using a VIAFLO 96 Electronic pipette. The plates were subsequently sealed with parafilm and incubated for another 24 hours at 37 °C, which resulted in the worms being stained red. To decrease the stain background, 240 μL of M9/0.1% Tween-20 was added to each well and centrifuged for 30 s at 2200×g. After removing 200 μL of liquid and adding 150 μL of M9/0.1% Tween-20, 25 μL of the mixture was transferred to another white 96-well plate containing 300 μL of M9/0.1% Tween-20. The white plates were scanned using a Epson Perfection V850 Pro flat-bed scanner using the settings configured to positive film holder, 3600 DPI, and 24-bit color. The images were processed in GIMP version 2.10.18. Horizontal and vertical gridlines were added to separate each well, and HTML color codes #000000 and #FFC9AF were removed. Unsharp masking was applied with a radius of 10, effect of 10, and a threshold of 0.5. Hue saturation adjustments were made: yellow, blue, cyan, and green hues had their lightness set to 100 and saturation to -100, while red and magenta hues had their lightness set to -100 and saturation to 100. Finally, each well was exported as a separate .tiff image using LZW compression. The number of worms in each well were counted using WorMachine in MATLAB, with the pixel binarization threshold set to 30, the neighboring threshold set to 1, and the maximum object area set to 0.003% (Hakim et al. 2018).

### Z-factor Measurements

We calculated the Z-factor for each 96-well plate from our screen. We did this using the formula: Z = 1 - (3 × (standard deviation of positive controls + standard deviation of negative controls)) / |mean of positive controls - mean of negative controls|.

### Structural Similarity Measurements

To assess structural relationships among 34 anti-microsporidia active compounds, Atom Pair fingerprints (APfp) (Bajusz et al. 2015; O’Boyle and Sayle 2016) were generated from SMILES structures and analyzed using ChemMine Tools (Backman et al. 2011). Hierarchical clustering was performed using single linkage with a Tanimoto similarity matrix derived from APfp fingerprints. Compound arrangement was visualized through a distance matrix heatmap with Z-score normalized activity values.

### Continuous infection assays

Compounds at a final concentration of 60 μM were incubated with 20,000 *N. parisii* spores/μL in 200 μL of K-medium in a 24-well plate. After one hour, L1 stage N2 worms in 200 μL of K-medium were added to each well. Plates were then covered with a breathable adhesive porous film, placed inside a humidity box which was wrapped in parafilm, and incubated at 21 °C with shaking at 160 rpm for 4 days. After incubation, all samples were washed with M9/0.1% Tween-20 twice, fixed with acetone, stained with DY96 as described below.

### Pulse infection assays

Approximately 8,000 L1 stage worms were mixed with 50 million *N. parisii* spores and 5 μL of 10x OP50-1 and added to a 6-cm NGM plates. Plates were dried and then incubated for three hours at 21 °C. To remove undigested spores, the worms were washed twice with M9/0.1% Tween-20. Plates were then set up as described above for the continuous infection assays, but without the addition of spores. Plates were incubated for either 2 or 4 days and then worms were fixed in acetone, and stained with FISH probe and DY96 as described below.

### Spore firing assays

*N. parisii* at a concentration of 20,000 spores/μl were incubated with 120 μM compounds and 2% DMSO for 24 hours at 21 °C. After washing the spores three times with K-medium, spores were used in the continuous infection assays as described above, without the addition of compounds and incubated for 3 hours. To determine if compounds could induce spore germination the spores were incubated with compounds for 24 hours as described in this section. Samples were then fixed in acetone and stained with a FISH probe and DY96 as described below.

### Mortality assay

*N. parisii* at a concentration of 20,000 spores/μL were incubated with 200 μM compounds and 2% DMSO for 24 hours at 21 °C. Acetone-treated spores were incubated with acetone for 24 hours. After incubation, spores were washed twice with 1 ml H2O, resuspended in 100 μL of 2 mg/L Calcofluor White M2R and 8 μM Sytox Green, and incubated at ∼21 °C for 10 minutes. Then, 2.5 μL of each mixture was spotted onto glass slides, mixed with 2.5 μL of 2% agar, covered with a glass slide, and examined using fluorescence microscopy as described below.

### DY96 staining, fluorescence *in situ* hybridization (FISH), and fluorescence microscopy

Samples were washed with M9/0.1% Tween-20 until most of the OP50 was removed. They were then fixed in 700 µL of acetone for 15 minutes and washed twice with PBS/0.1% Tween-20. The samples were then rotated for 30 minutes in a DY96 solution (10 µg/mL DY96, 0.1% SDS in 1x PBS + 0.1% Tween-20), after which EverBrite mounting medium with DAPI was added. FISH staining was carried out using the microB FISH probe for *N. parisii* 18S rRNA (ctctcggcactcctcctg) conjugated to Cal Fluor Red 610 (LGC Biosearch Technologies) (Troemel et al. 2008). After being washed with PBS/0.1% Tween-20 and hybridization buffer (900 mM NaCl, 20 mM Tris HCl, 0.01% SDS), samples were incubated overnight at 46 °C with the hybridization buffer containing 5 ng/mL FISH probe. Subsequently, the samples were washed once with 1 mL of hybridization buffer containing 5 mM EDTA. The samples were stained with DY96 as previously described above and mounted on glass slides with EverBrite. Images of the samples were obtained using Zeiss Zen 2.3 software with a ZEISS Axio Imager 2 microscope at magnifications ranging from 5x to 63x. Worms containing any number of embryos were classified as gravid. An infected worm was defined as an animal exhibiting any newly formed spores.

### *P. epiphaga* infection assays

Stocks of *P. epiphaga* (JUm1396) spores were prepared as previously described (Murareanu et al. 2022). Briefly infected populations of N2 worms were cultured on NGM plates seeded with OP50. Infected worm populations were then processed as described above for *N. parisii* stocks. The infection experiments with *P. epiphaga* followed the same procedure as the continuous infection assays. Each well contained a final volume of 400 μL of K-medium, comprising 800 L1 worms, 80,000 *P. epiphaga* spores/μL, and either 60 μM of the compound or 1% DMSO. Fixing the samples in acetone and washing twice was followed by staining with the FISH probe specific to *P. epiphaga* 18S rRNA (CAL Fluor Red 610CTCTATACTGTGCGCACGG). FISH fluorescence was quantified under fluorescence microscopy.

## Statistical analyses

Data from each experiment consisted of three independent experiments biological replicate and analyzed using GraphPad Prism. P-values were determined by ANOVA, with statistical significance defined as follows: p<0.05, *p<0.01, ***p<0.001, and ****p<0.0001.

## Supplementary materials

**Fig. S1.**
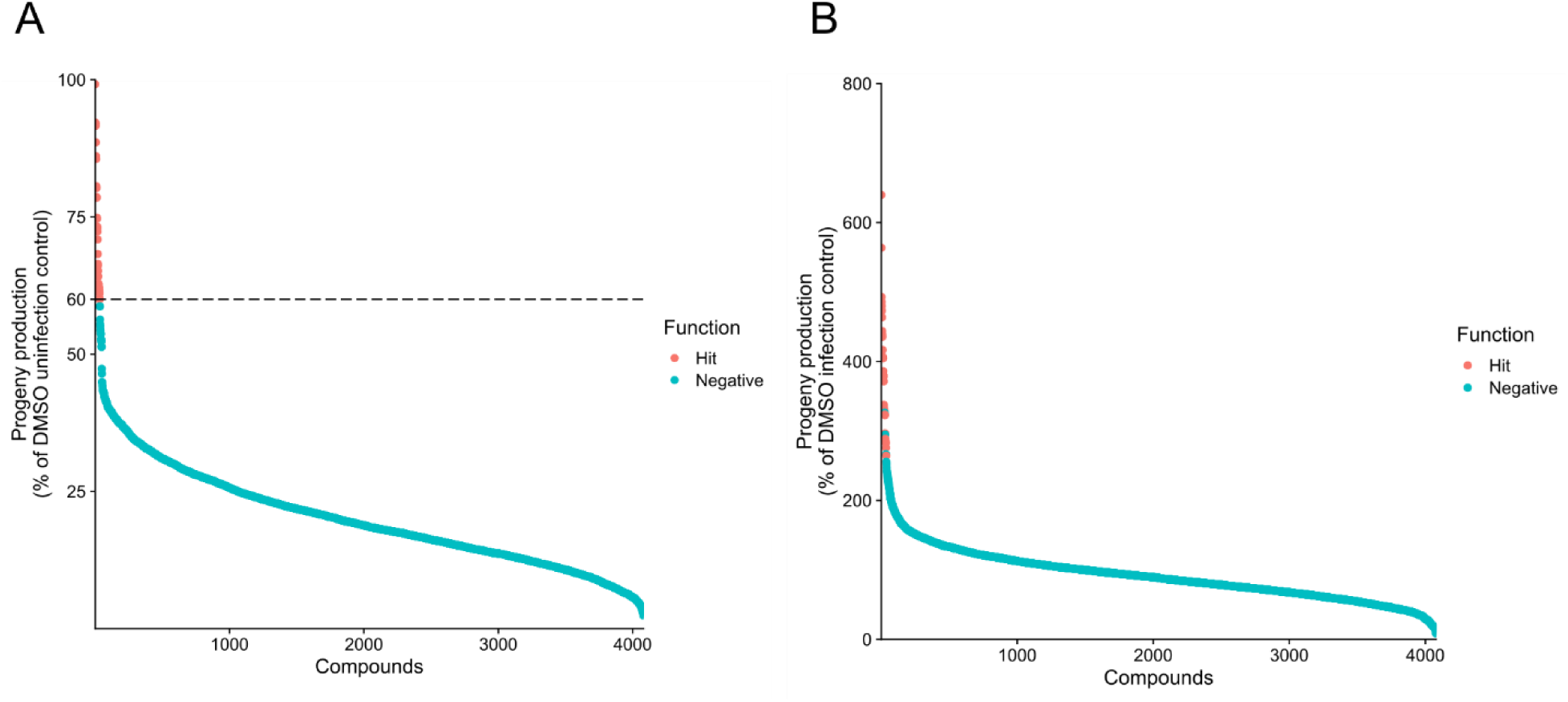
Ranked progeny production and fold increase for 4,080 BU-CMD compounds tested against *N. parisii* infected *C. elegans.* **(A)** Data described in figure 1A, which is the percentage of progeny production compared to uninfected controls, is presented in ranked order. **(B)** The percentage of progeny production compared to infected controls. In both (A) and (B) the compounds are colored as in Figure 1A with compounds with progeny production less than 60% are shown colored blue and compounds with progeny production of at least 60% are shown colored red.

**Figure S2.**
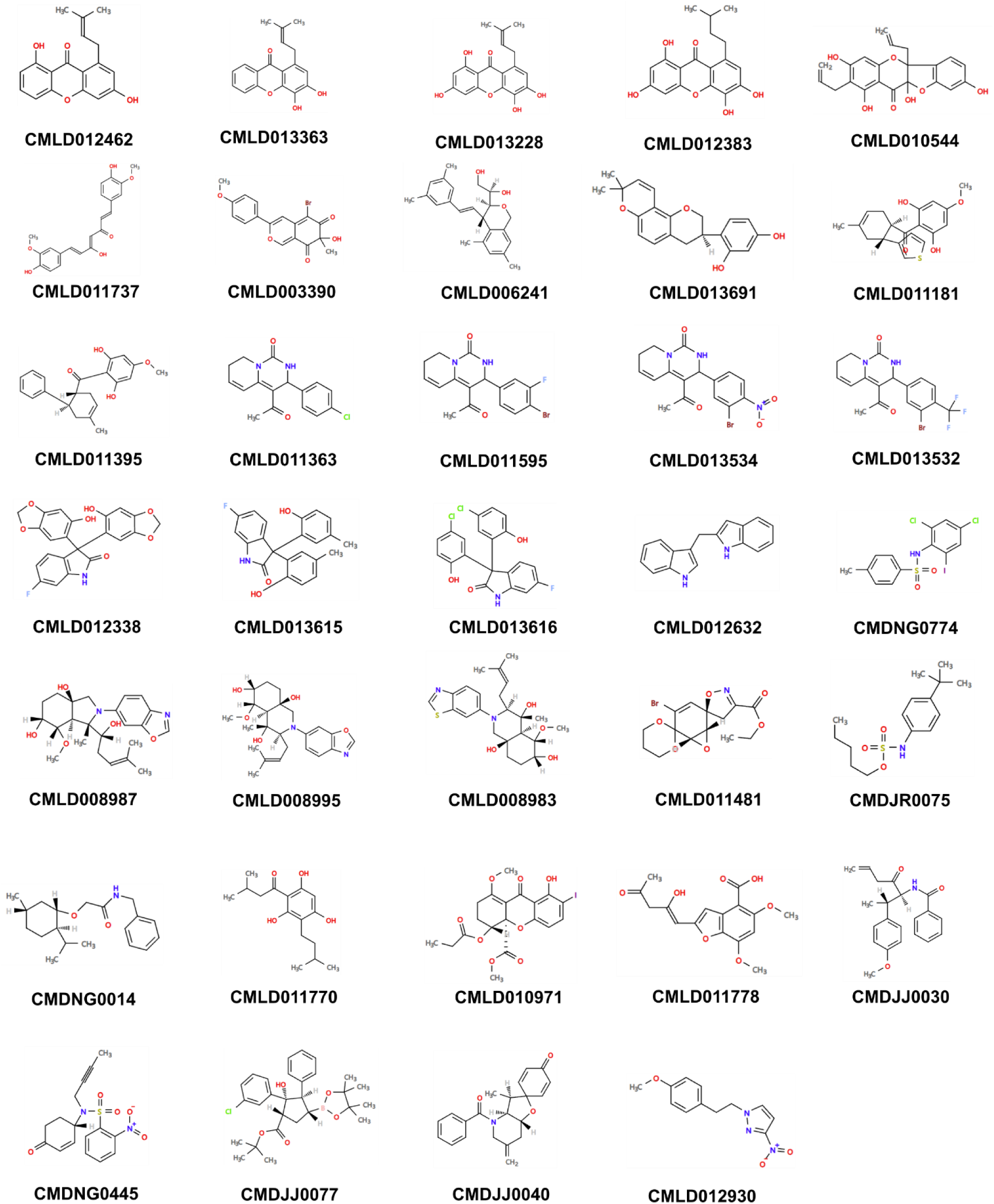
Chemical structures of the 34 BU-CMD compounds with progeny production of at least 60%. The structure of each compound is shown with the label below the compound. Compounds are ordered the same as in Figure 1B.

**Figure S3.**
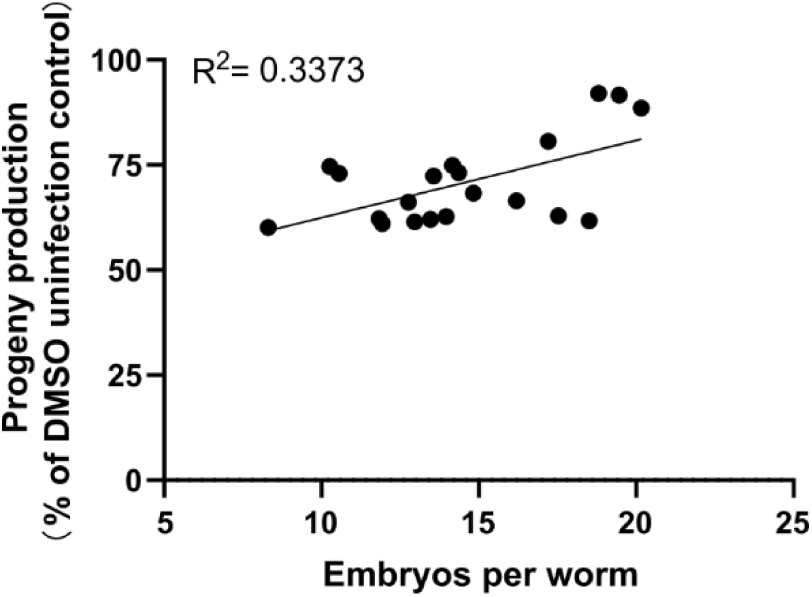
Progeny production from a screen of BU-CMD compounds is modestly correlated with number of embryos per worm from validation experiments. A linear correlation was performed between the percentage progeny production from Figure 1A and the number of embryos per worm for the 20 BU-CMD compounds tested in Figure 2A.

**Figure S4.**
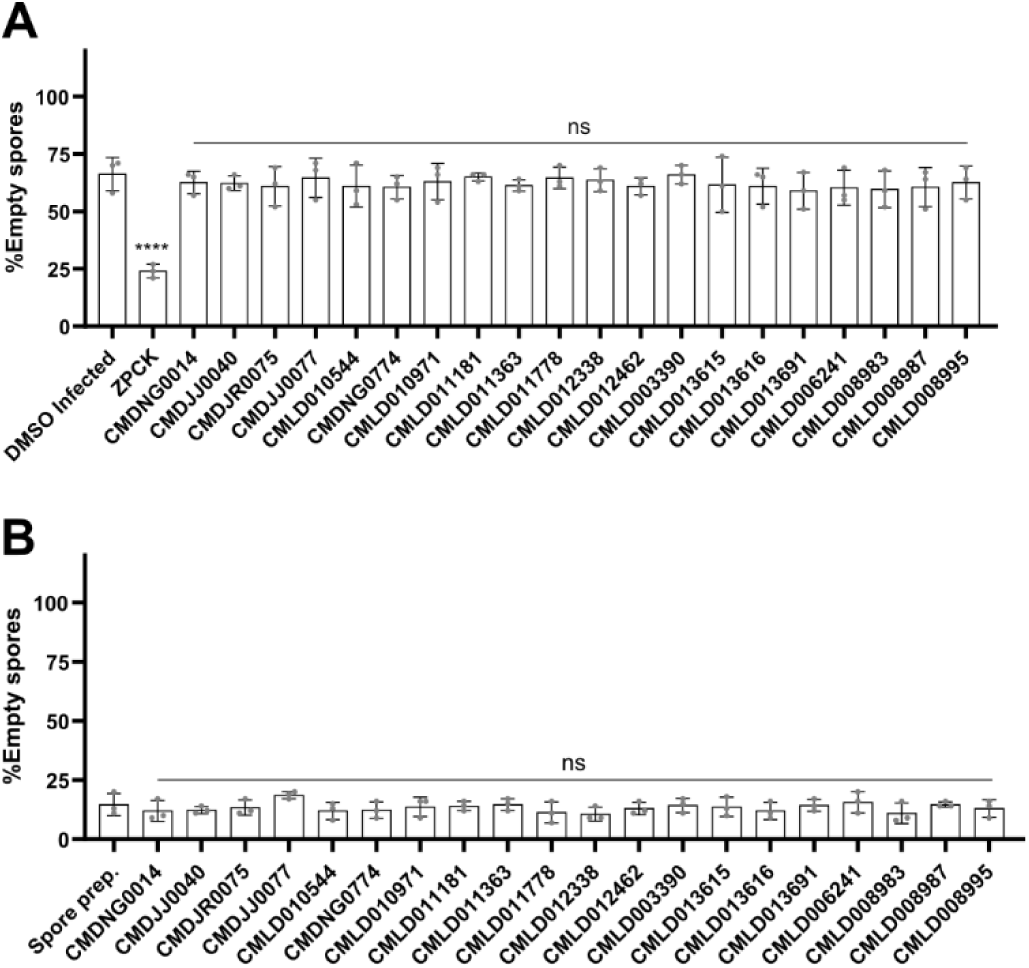
Validated BU-CMD compounds do not prevent spore germination in vivo nor trigger sporing firing *in vitro*. **(A)** *N. parisii* spores were incubated with compounds for 24 hours and the compounds were washed away from the spores. These spores were then incubated with L1 stage worms for 3 hours, and then worms were fixed and stained with FISH probe specific to the *N. parisii* 18S rRNA and DY96. Percentage of spores not containing a sporoplasm was quantified. **(B)** *N. parisii* spores were incubated with compounds for 24 hours and the compounds were washed away from the spores. Spores were then fixed and stained with FISH probe specific to the *N. parisii* 18S rRNA and DY96. Percentage of empty spores (spores not containing a sporoplasm). n = 3, N = ≥ 100 spores counted per biological replicate. The P-values were determined by one-way ANOVA with post hoc test (***p < 0.001 and ns means not significant). Means ± SD (horizontal bars) are shown.

